# chroGPS2: differential analysis of epigenome maps in R

**DOI:** 10.1101/425892

**Authors:** Oscar Reina, Fernando Azorin, Camille Stephan-Otto Attolini

**Affiliations:** Institute for Research in Biomedicine (IRB Barcelona), The Barcelona Institute of Science and Technology, Baldiri Reixac, 10, 08028 Barcelona, Spain; Institute of Molecular Biology of Barcelona, CSIC, Baldiri Reixac, 10, 08028 Barcelona, Spain

**Keywords:** Chromatin, Epigenetics, Visualization, Functional analysis, Differential analysis, Multidimensional Scaling, Clustering

## Abstract

1

**Backgrounds:** In the last years we have faced an unprecedented growth in the availability of high-throughput epigenomics data related to the genomic distribution of epigenetic marks, namely transcription factors, histone modifications and other DNA binding proteins. This also pointed out the need for efficient tools for integration, visualization and functional analysis of genomics and epigenomics data. On this subject, we previously developed a computational framework, chroGPS, to integrate and visualize the associations between epigenetic factors and their relation to functional genetic elements in low-dimensional maps. We demonstrated the usefulness of our approach with several practical case studies based on well-defined biological hypothesis.

**Results:** Here we introduce chroGPS version 2, a major update of our previously developed software with new functionalities for differential analysis of epigenomes. Methods are provided for efficient integration and comparison of data from different conditions or biological backgrounds, accounting and adjusting for systematic biases in order to provide an efficient and statistically robust base for differential analysis. We also include new useful functionalities for general data assessment and quality control prior to comparing maps, such as functions to study chromatin domain conservation between epigenomic backgrounds, to detect gross technical outliers and also to help in the selection of candidate marks for de-novo epigenome mapping. Our software, implemented in R as a Bioconductor package, provides detailed reference and user manuals to allow the final user to reproduce the presented case studies and use them as a starting point for generating and exploring epigenetic maps according to their own experimental needs.

**Conclusion:** chroGPS2 extends our previously developed software, providing now a complete, intuitive, efficient and statistically robust framework for generation, visualization, characterization and differential analysis of epigenomic maps, in a way which is easy to adapt to different biological and technical backgrounds allowing exploration and hypothesis testing in multiple biological scenarios.

## 3 Background

In the last years, the availability of epigenomics data related to whole-genome distribution of transcription factors, histone modifications and other DNA binding proteins has increased exponentially [Celniker <u>et al.</u>, 2009; Dunham <u>et al.</u>, 2012; Bernstein <u>et al.</u>, 2010], helping to establish a deeper knowledge about chromatin states and topologically associated domains [Filion <u>et al.</u>, 2010; Serra <u>et al.</u>, 2016], and to offer a broad scope of genomic regulation from both a functional and structural point of view. However, there is still a clear need for efficient tools for visualization, functional analysis and comparison of epigenomics data [Marx, 2015]. Previously, we developed chroGPS [Font-Burgada <u>et al.</u>, 2014], an R package based on dimensionality reduction techniques (namely Multidimensional Scaling, MDS) to measure and visualize genome-wide associations between epigenetic factors and their relationship to functional genetic elements in intuitive and easy to interpret two or three-dimensional maps. Now we extend this software and introduce novel features to perform differential analysis of epigenome maps using Procrustes [Lisboa <u>et al.</u>, 2014], hierarchical clustering and Bayesian density estimation methods [Jara <u>et al.</u>, 2011]. Additionally, we provide new functions for general data assessment, quality control and to help in the selection of epigenetic factors for de-novo epigenome mapping.

## 4 Implementation

ChroGPS2 is integrated in Bioconductor [Huber <u>et al.</u>, 2015], an open-source collection of R packages for computational analysis of omics data. Essentially, the core methodology used to generate chroGPS maps as described in detail in [Font-Burgada <u>et al.</u>, 2014] starts by measuring similarities between epigenetic factors or genomic features based on their whole genome binding profile overlaps (chroGPS-factors and chroGPS-genes respectively), using the function *distGPS*. After this, two or three-dimensional maps for visualization are generated using the *mds* function, which provides both metric and non-metric MDS variants, as well as additional optimization steps and a parallel implementation developed in order to speed up generation of chroGPS-genes maps, which can involve tens of thousands of points. Finally, the *clusGPS* function can be used to perform functional analysis of the generated maps using hierarchical clustering. We now introduce new functionalities to perform additional quality control and data assessment steps prior to generation of epigenomic maps. The *domainDist* function does that by assessing the degree of conservation between chromatin domains defined by certain groups of epigenetic factors, and can be used to assess presence of gross technical problems or annotation errors present in the data that could affect subsequent integration or differential analysis. Additionally, the *rankFactors* functions can help researchers with selection of candidate marks when designing de-novo epigenome mapping experiments in order to compare a novel background or condition with already existing ones, a scenario posing important technical and economical challenges. Differential analysis of chroGPS-factor maps is performed using Procrustes, and relevant differences between groups are assessed via Procrustes sum of squared errors using the *diffFactors* function. Differential analysis of chroGPS-genes is performed using hierarchical clustering and Bayesian density estimation methods with the *diffGenes* function. Additional helper functions are also provided to integrate data from different biological or technical background, to deal with unification of biological or technical replicates, and to export the generated maps in interactive HTML format.

## 5 Results

### 5.1 Data assessment and Quality Control

We offer functions to perform a first general assessment of functional relationships between epigenetic factors and conformation of chromatin domains across datasets, and to highlight major differences between them. After using the *distGPS* function to measure pairwise similarities between epigenetic factors based on genome-wide overlap of their binding profiles, the *domainDist* function evaluates and compares these results at factor and domain level, pointing towards major changes that may suggest strong biological effects but also gross technical problems or annotation errors (i.e. sample mislabeling, truncated or empty data, inefficient immunoprecipitation, see Supplementary section 2.1).

### 5.2 Select candidates for de-novo epigenome experiments

Selection of candidate factors when designing high-throughput ChIP-Seq, Dam-ID or Hi-C experiments in order to compare a known scenario with a novel one poses important experimental and economical challenges. The *rankFactors* function effectively ranks a set of factors based on how much they contribute towards conservation of their respective chromatin domain identity. Additionally, if domain identity is unknown or incomplete we rely on relative conservation of functional relationships between individual epigenomic marks and genetic elements (i.e. genes, promoters, etc) and how information from experimentally mapped factors can be used to successfully impute presence/absence of others [Ernst and Kellis, 2015]. In detail, we offer methods based on linear and logistic regression to rank marks based on their ability to predict others, helping the researcher to refine selection of potential candidates for experimental mapping (See Supplementary section 2.2).

### 5.3 Comparing epigenomic factor maps

chroGPS factor maps use MDS to represent genome-wide epigenetic factor co-localization based on similarities between their binding profile overlaps according to a metric of choice. This core methodology is also used as a starting point to identify differences between epigenomic factor associations at different conditions, time points, cell lines, or between different species.

Comparing two epigenomic factor maps from the same species with a rich number of common factors is relatively straightforward, as they share a genomic background over which pairwise similarities between their genome-wide binding profile overlaps can be computed using the *distGPS* function, and represented together in a low dimensional space using MDS. This already produces a joint map which accurately represents putative functional relationships between all elements from both datasets [Font-Burgada <u>et al.</u>, 2014]. Further adjustment and differential analysis is performed with the function *diffFactors*, which uses Procrustes to identify potential biological differences between both backgrounds after adjusting for possible technical and biological biases. Procrustes matches two sets of points represented in a low-dimensional space by using the information from common elements (landmarks) to compute a transformation involving scaling, translation and rotation to minimize euclidean distances between landmark points while preserving relative spatial configuration within each set. General goodness of fit and magnitude of the observed changes between common factors is measured via Procrustes sum of squares, and statistical assessment of the changes in epigenomic overlap can be performed via permutation tests [Gel <u>et al.</u>, 2016]. This approach has been successfully applied to integrate and compare the epigenomic landscape obtained at different time points during genome replication in *Drosophila* S2 cells (See Figure 1 and supplementary section 3.3).

**Figure 1:**
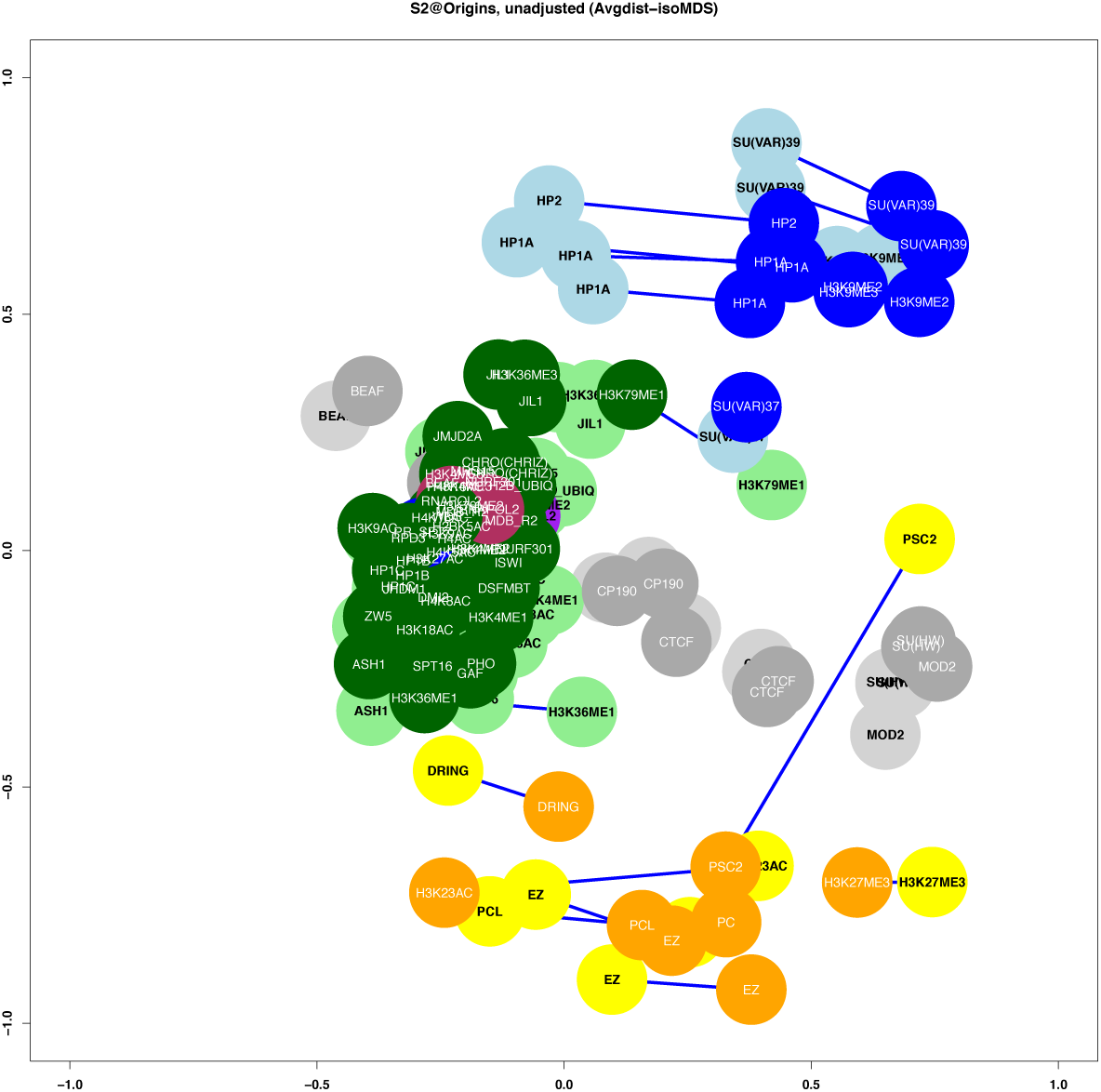
Differential chroGPS-factors map showing the epigenomic landscape differences at *Drosophila melanogaster* Early (light) and Late (dark) time points during genome replication

### 5.4 Comparing epigenomic gene maps

ChroGPS-genes maps represent similarities between genomic features by computing pairwise distances between the binary vectors defined by their epigenetic footprint, that is, their collection of nearby epigenetic marks, and projecting them in a low dimensional space using MDS [Font-Burgada <u>et al.</u>, 2014]. Downstream functional analysis can be performed by using hierarchical clustering with the *clusGPS* function or using the original and approximated distances with other machine learning or class discovery methods of choice.

- **Detecting epigenomic footprint differences:** The *diffGenes* function uses this strategy to compare two sets of mapped factors from the same or different species using epigenomic footprint information of both datasets simultaneously. First, common factors between both sets are selected. Then, pairwise similarities between their whole-genome epigenetic footprints are computed to generate a map comprising all common genes from both datasets, relying on inter-species gene homology information if necessary. Identification of genes presenting changes in their epigenomic footprints is then straightforward, as they will be indicated by those genes being represented by different points for each set.
- **Interpreting loss of epigenomic footprint identity:** A first approach to study those differences is to rank them using Procrustes to compute the residual sum of squares between points representing differentially located genes, but this offers little insight on the observed changes from a functional perspective. We use unsupervised hierarchical clustering followed by computation of Bayesian posterior probability of cluster classification using a Dirichlet Process mixture of normals [Jara <u>et al.</u>, 2011] to identify and characterize clusters of genes with similar epigenetic patterns, and therefore to detect genes classified under distinct clusters between both conditions. Akin with results obtained from other high-throughput technologies, candidate gene lists can be used for further downstream functional analysis, such as Gene Ontology enrichment or Gene Set Enrichment Analysis, as well as compared with other genomics data such as changes in gene expression or regulation to further explore interesting hypothesis. We illustrate the use of this strategy to identify and characterize genes potentially involved in epigenomic changes between third instar larvae *Drosophila* S2 cells and BG3 neuronal tissue. (See Figure 2 and Supplementary section 4.3).

**Figure 2:**
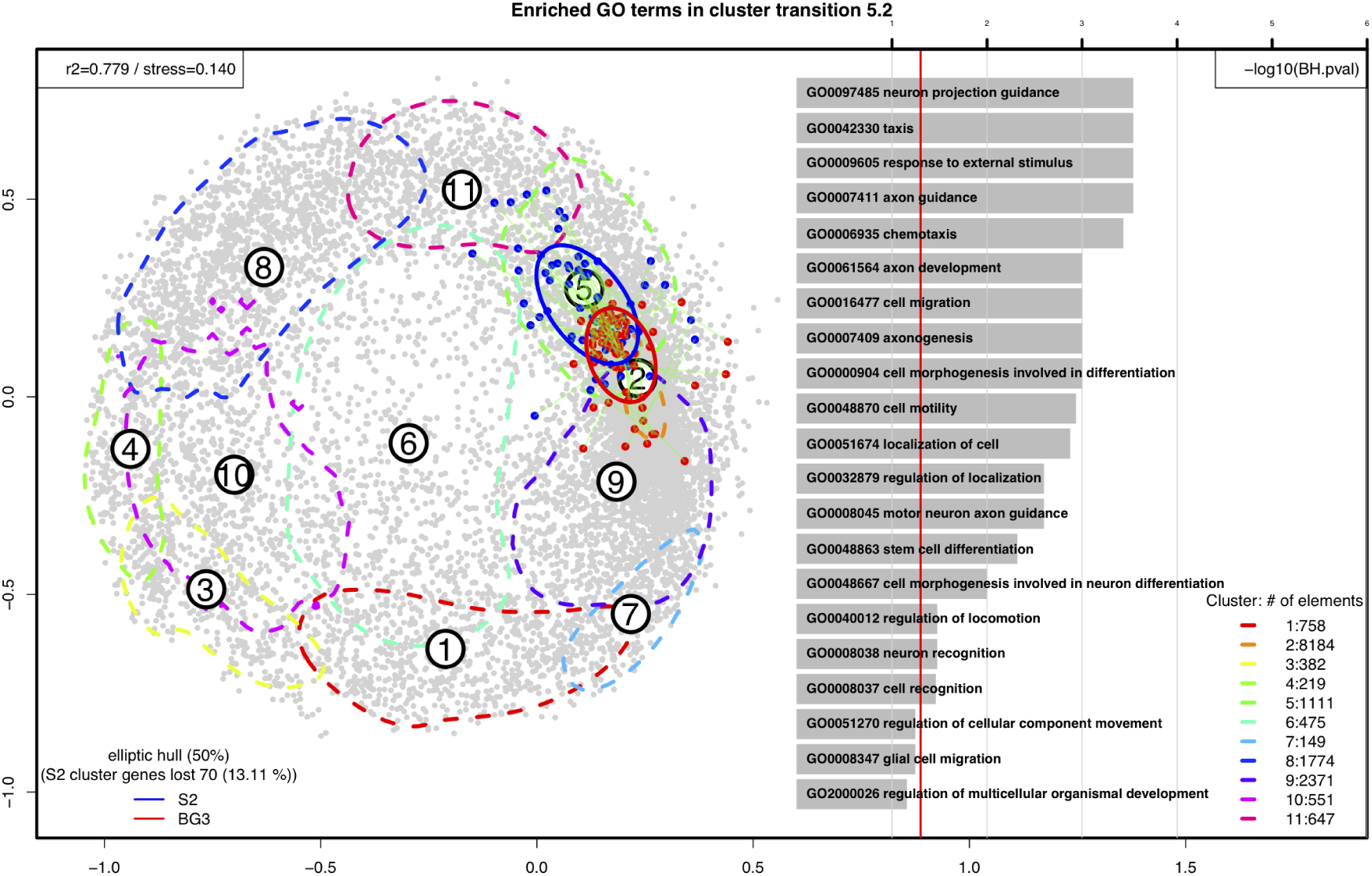
Differential chroGPS-genes map of *Drosophila melanogaster* S2 and BG3 cell lines. Focus is put on genes changing from cluster 5 in S2 (blue, moderate HP1 repression) to cluster 2 in BG3 (red, transcriptionally active). Dashed lines indicate probability contours containing 50 percent of genes for each cluster. On the right, the top 20 enriched Gene Ontology terms for genes involved in the analyzed cluster transition are shown. Selected genes are specially enriched in nervous system and cell differentiation categories.

## 6 Conclusions

Here we presented chroGPS2, a new major version of our Bioconductor package chroGPS, for visualizing and performing functional and differential analysis of epigenomics data using dimensionality reduction techniques (Multidimensional Scaling, MDS). Usefulness of our approach is illustrated using several examples to generate and compare 2-dimensional maps from publicly available ChIP-chip and ChIP-Seq fruit fly data, that can be also applied in order to incorporate other kinds of data used to study chromatin conformation such as Hi-C or DRIP-Seq. Methods are provided for integration of genomics and epigenomics data from different technical or biological backgrounds, accounting and adjusting for systematic biases in order to provide an efficient and statistically robust base for differential analysis. We also offer fast and user-friendly computational solutions for visualization and differential analysis to assess changes between compared groups, conditions or species. Along with the case-studies introduced here, we provide extensive supplementary material with additional information and a detailed package vignette with reproducible code including step-by-step instructions to generate and compare epigenomic maps from public data available in modENCODE, and that can be used as a guideline to explore and test novel hypotheses using publicly available and in-house generated data in a wide range of biological scenarios.

## 7 Availability and requirements

Project name: chroGPS2

Project URL: https://www.bioconductor.org/packages/release/bioc/html/chroGPS.html

Operating systems: Linux, Mac OS X, Windows

Programming language: R language

Other requirements: R 3.2.0 or higher. Required packages: GenomicRanges, IRanges, methods, Biobase, MASS, graphics, stats, changepoint, cluster, DPpackage, ICSNP, ellipse, vegan

License: GPL 2.14

