## Supplementary material for "chroGPS2: differential analysis of epigenome maps in R"

### chroGPS2: differential analysis of epigenome maps in R.
Supplementary material

Oscar Reina[1], Fernando Azorin[1,2] and Camille Stephan-Otto Attolini[1].

[1]Institute for Research in Biomedicine (IRB Barcelona), The Barcelona Institute of Science and Technology, Baldiri Reixac, 10, 08028 Barcelona, Spain.
[2]Institute of Molecular Biology of Barcelona, CSIC, Baldiri Reixac, 10, 08028 Barcelona, Spain.

May 14, 2019

**6 DATA AND CODE AVAILABILITY** **14**

### 1 DATA ACQUISITION AND FORMATTING

The source material for generating and comparing epigenomic maps with chroGPS is a list of genome-wide predicted binding sites for each condition, formatted as a *GRangesList* object, that is, a list of Genomic Ranges. These can simply be collections of plain-text BED or GFF files downloaded from ENCODE, modENCODE, Roadmap Epigenomics [Bernstein et al., 2010; Celniker et al., 2009; Dunham et al., 2012], or any other public data repository, or in-house generated data coming from any of the widely used Peak Calling algorithms such as MACS, Sicer, etc. Such files can be easily imported into R using custom code or readily available functions from other Bioconductor packages [Gentleman et al., 2004], as for instance, *rtracklayer*.

### 2 DATA EXPLORATION AND QUALITY CONTROL

Before performing differential analysis some useful information can be obtained from original data to determine goodness of the selected datasets and help interpreting the observed differences afterwards. In detail, we offer functions to study the degree of conservation between chromatin domains defined by groups of factors under two given conditions, and to assess presence of gross technical problems or annotation errors present in the data that could affect subsequent integration and comparison. Additionally, we implement methods to help researchers with selection of candidate marks when designing de-novo epigenome mapping experiments in order to compare a novel background or condition with already existing ones, a scenario posing important technical and economical challenges.

#### 2.1 Exploring factors and domains

The core methodology of chroGPS, that is, computation of pairwise similarities (and thus, dis-similarities that can be interpreted as distances) between epigenomic factors based on their binding profile overlaps, already provides some useful insight on relative configuration of epigenomic factor domains present in the data. In detail, this information can be compared across multiple datasets from which a rich number of common mapped factors is available, to assess correlation in vectors of similarities between pairs of analog factors or domains, in order to identify potentially strong biological or technical differences between them. We make use of this functionality using the *domainDist* function to assess the already observed general domain conservation between *Drosophila melanogaster* S2 and BG3 cell lines [Celniker et al., 2009; Font-Burgada et al., 2014], and to see what happens when we artificially introduce a 'wrong' instance of factor EZ in the S2 dataset (See Supplementary Figure 1).

We will start by loading an object with downloaded Binding Sites for several *Drosophila melanogaster* S2 and BG3 epigenomic factors annotated for closest and overlapping dm3 genes using the *AnnotatePeakInBatch* function from the *ChIPpeakAnno* package [Zhu et al., 2010], and will unify experimental replicates by simply joining all reported binding sites for each factor. Additional methods of replicate management are provided in the function *mergeReplicates*.

Afterwards, we will use the color information provided in the *s2names* data frame object, which will effectively serve as aliases for our chromatin domains of interest. And then, we will take care of computing pairwise similarities / distances between S2 and BG3 epigenomic factors based on their whole genome binding profile overlaps [Font-Burgada et al., 2014], and will use the *domainDist* function to calculate both within and between-domain distances, which can be described as the collection of computed mathematical distances measured between all pairs of factors belonging to a certain chromatin domain (intra), or between all possible pairs of factors from two different domains (inter). This information gives a valuable insight regarding cohesion and separation of the different domains observed in the map, and can be used to start assessing putative conservation between factors on different biological backgrounds.

The returned intra and inter-domain distance objects can be used for generating custom plots and performing further downstream analysis, and the observed differences between datasets can be assessed statistically via permutation tests [Gel et al., 2016]. We can also see the effects of introducing an artificial outlier EZ sample in the S2 cell line, by randomly selecting binding sites from other factors.

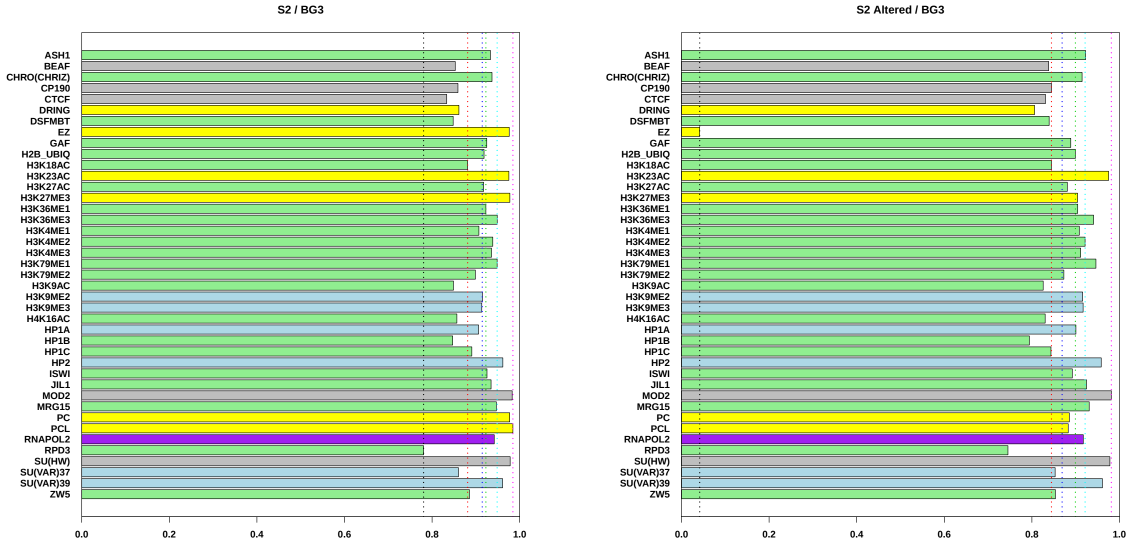

Supplementary Figure 1: Detection of potential technical artifacts by checking conservation of relative distances between common factors. Left: Conservation for *Drosophila* S2 and BG3 epigenomic datasets. Right: The same plot where artificially modified EZ factor is introduced in the S2 dataset.

#### 2.2 Selecting factors for de-novo epigenome mapping

**Maximize chromatin domain identity:** Chromatin domains offer an insightful and intuitive way to interpret epigenomic map conformation, by providing a biological context to factors based on functional relationships between them. When such information is available, a straightforward approach to select candidate factors for performing a de-novo epigenome mapping is to select those ones giving maximum robustness to their corresponding domain. This is easily achieved using the *rankFactorsbyDomain* function. We can use our chromatin color values as an alias to define chromatin domains. In this example we see how to perform a domain distance based selection for the HP1a repression considering a subset of 4 different factors (See Supplementary Figure 2).

**Ranking factors based on functional relationship with genetic elements**. Alternatively, or whenever domain information is not available, selection of candidate factors can be based upon conservation of functional relationships between epigenetic factors and genetic elements and how information on experimentally mapped factors has been successfully used to impute unknown ones [Ernst and Kellis, 2015]. The data for performing this analysis is that one used to generate chroGPS-genes maps (See Supplementary section 4), that is, a binary matrix of $N$ genes (rows) per $M$ columns (factors), where each cell equals 1 if that gene has an assigned binding site for that factor, and 0 otherwise. The *rankFactorsbyProfile* function also provides methods based on linear and logistic regression to rank factors based on how accurately they can be predicted by others. At each iteration, the factor which can be best predicted by the rest is removed and prediction accuracies are recomputed (see Supplementary Figure 2). On the downside, these methods can be computationally intensive and we recommend using parallel computation or reducing the number of iterations if computing power is limited.

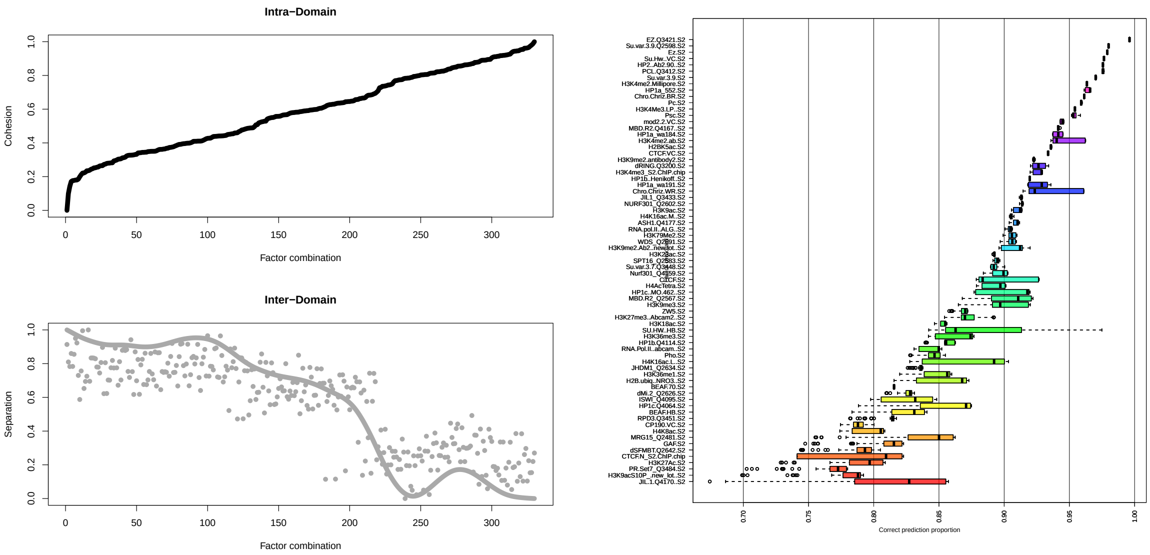

Supplementary Figure 2: Left: From left to right, resulting domain integrity (Cohesion/Separation) based on computation of Intra and Inter-Domain distances for each combination of 4 factors within the HP1a repression domain. Results are sorted by Intra-Domain distance, which is an informative concept regarding cohesion of the observed domains in the map. Potential candidates are those among the leftmost factor combinations minimizing Intra-Domain distance but still providing meaningful biological content. Right: Factor ranking based on logistic regression of functional relationship with genetic elements when no domain information is available. Left axis, from top to bottom, indicates factor removed at each iteration, which is the one having the higher correct prediction rate based on information from those ones remaining below, until the number of remaining factors is lower than a given threshold. Boxplots in each line indicate the obtained prediction rates for all removed factors on top, based on the information from those on the bottom. Notice that as we remove more factors, general prediction rates get lower.

### 3 COMPARING CHROGPS-FACTORS MAPS

In chroGPS-factors maps epigenetic factors are represented over a low-dimensional space based on a similarity measure based on their observed co-occurences at whole genome level. However, this is only a global picture of epigenetic factor colocalization and thus does not reflect the possible mechanisms going on in certain kind of genomic regions (i.e. coding regions, promoters, transcription start sites, enhancers etc). Epigenomic factor colocalization is indeed a complex scenario, playing a critical role in gene regulation and silencing, transcriptional replication, genome structure and repair, etc. A very straight-forward exercise is to observe distribution of these epigenetic marks based on factor overlaps happening only over certain specified regions of interest, to be used at subsequent comparisons. Even though the methodology to generate chroGPS-factors maps is available in our previous work, first we will provide a brief reminder on how these maps are generated.

##### 3.1 Generating factor maps

The main ingredient for generating chroGPS-factors maps is a collection of genomic intervals from our epigenomics data (putative binding sites, enriched regions, etc) in *GenomicRanges* format, in the shape of a *GRangesList* object, belonging to the experimental condition we want to study (i.e. cell line, patient, etc). Such information can be stored in independent plain tab-separated, BED or GFF files. Once our genomic intervals collection is loaded, we make use of the distGPS function to compute pairwise similarities / distances between them, as a way to assess whole genomic co-localization in our epigenomics background. At chroGPS we provide different metrics designed to work with genomic interval data to account for several scenarios where certain mathematical characteristics may be desirable, as well as the option to feed the workflow with user-defined metrics [Font-Burgada et al., 2014]. The resultant object contains a *n x n* matrix of dissimilarities ranging between 0 and 1 that can be already used for visualization and analytical purposes (i.e. heatmaps, clustering).

The next step is to use the distance object as input to the core of our methodology. Multidimensional Scaling (MDS) technique used used to dissimilarity data in easy to visualize, easy to interpret, 2 or 3 dimensional graphical representations that account for the original relationships of similarity between the observed objects [Borg and Groenen, 2003]. We account for several MDS methods, and provide functions for parallel computation and goodness-of-fit optimization when several thousands of elements are present in our dataset.

The returned object can be represented in a 2 or 3 dimensional space where each element is located based on their similarity (i.e. co-occurrence) with all the other elements at whole genome level. This strategy already proved successful at reflecting the biological nature of epigenomic domains in *Drosophila melanogaster* S2 cells as illustrated in the left panel of Supplementary Figure 3. MDS objects returned can be plotted directly using the package provided method. Additionally, the function *getPoints* can be used to retrieve values for each element present in the map and used for producing custom or enhanced graphical outputs.

##### 3.2 Generating region-specific factor maps (Promoters)

The basis for this case study is the same epigenetic factor collection used to generate the *Drosophila melanogaster* map above, plus additional information providing the genomic regions in which we would like to focus. In our case we will focus our study over gene promoters by obtaining UCSC dm3 genes and generating a new *GenomicRanges* collection with promoter regions (defined as 1kb upstream of the TSS).When these two objects are provided as input for the *distGPS* function, similarities between epigenetic factors are only taken into

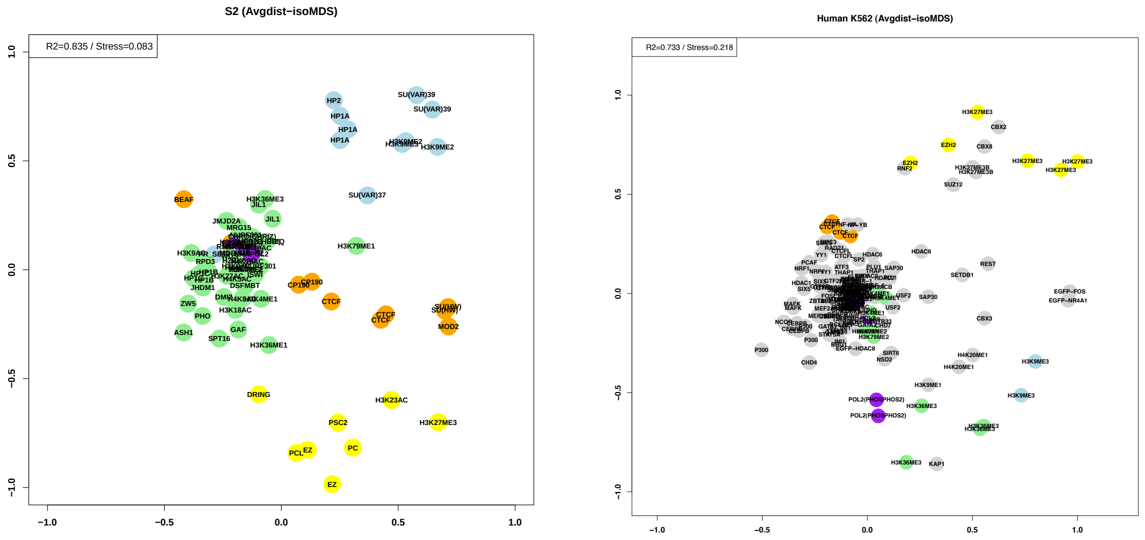

Supplementary Figure 3: chroGPS-factors map of *Drosophila melanogaster* S2 cells (left) and human K562 leukemia cell line (right).

account when their overlap falls over the specified regions of interest. Thus, co-occurrences taking place at coding or intergenic regions will be ignored. If more control over region-based filtering is needed, interval-based operations from the IRanges and *GenomicRanges* packages offer advanced options to address it.

Straightforward visualization of the map generated in this way (See Supplementary Figure 4) already provides some insights on potential regulatory changes happening at promoter regions. Direct visual comparison with the whole genome *Drosophila melanogaster* S2 map [Font-Burgada et al., 2014] already highlights interesting changes in the conformation of some epigenomic domains, such as location of the Heterochromatin Protein complexes. However, it is desirable to perform further analitical work to offer the user quick and unbiased identification of significant changes when comparing region-based maps between two collection of regions or against a whole-genome scenario.

#### 3.3 Comparing factor maps between conditions (Origins of Replication)

Procrustes allows integration of epigenetic maps coming from different biological backgrounds, as well as adjustment of undesired biases due to technical effects [Borg and Groenen, 2003; Kendall, 1989]. Furthermore, it can be used to identify differences between epigenomic factor maps generated at different conditions (healthy/disease, control/treated), coming from distinct biological backgrounds, or being snapshots of different points in time, such as developmental stages or timings in origins of replication. We will focus on this elements to illustrate comparison of chroGPS-factors maps.
Origins of replication are particular genomic sequences at which replication of DNA starts in living eucharyote and prokaryote organisms, and of DNA-RNA in viruses. These sequences are recognized by specific proteins, that recognize, unwind and begin to copy the genomic

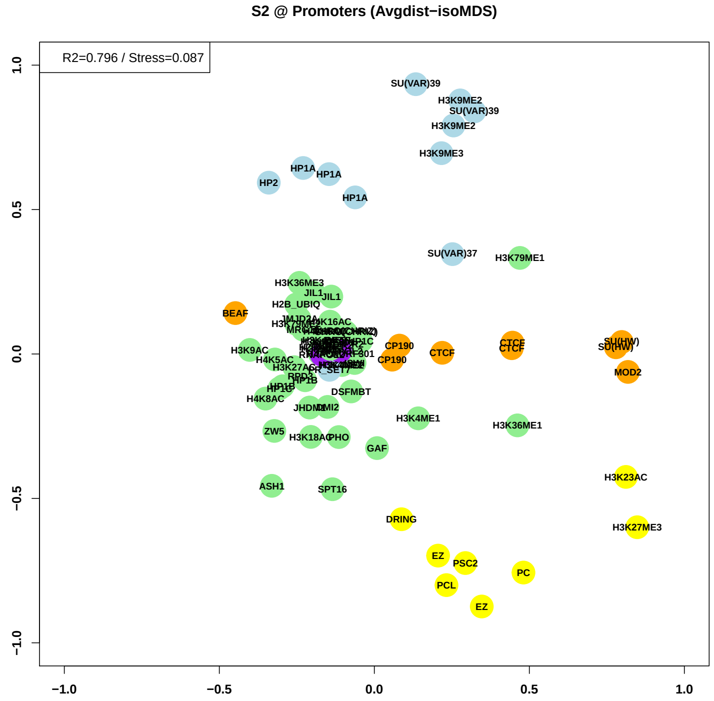

Supplementary Figure 4: chroGPS-factors map of factor co-localization over *Drosophila melanogaster* S2 promoters.

sequence. In the following steps we illustrate how to use chroGPS to compare the epigenomic landscape at these different time points.

First, we start by loading Origins of Replication data from *Drosophila melanogaster* S2 cells available in modENCODE [Celniker et al., 2009], and contains location for Origins of Replication at Early, Early Mid, Late Mid and Late cell replication time points, stored in four different BED files. The other data to perform our study will be our already familiar collection of epigenomic elements (genomic intervals) in all conditions we would like to compare. In our case we will use *Drosophila melanogaster* S2 modENCODE binding sites and will filter them according to the different collections of Origins of Replication regions we just loaded. This can be easily performed using basic IRanges overlap operations.

The next step is to generate regular chroGPS-factors maps of each joint dataset as described in section 3.1. Map comparison is performed using Procrustes to measure and rank changes between two given maps with the *diffFactors* function. This function performs the dual task of finding an optimal adjustment in shift, scale and rotation so that both maps are matched as much as possible while maintaining relative distances between the respective factors of each one and providing the user with the internal sum of error metrics of the Procrustes algorithm. As a result, we obtain both a graphical 2D chroGPS-factors map where distances between replicates of the same element in both backgrounds are highlighted, and a ranked

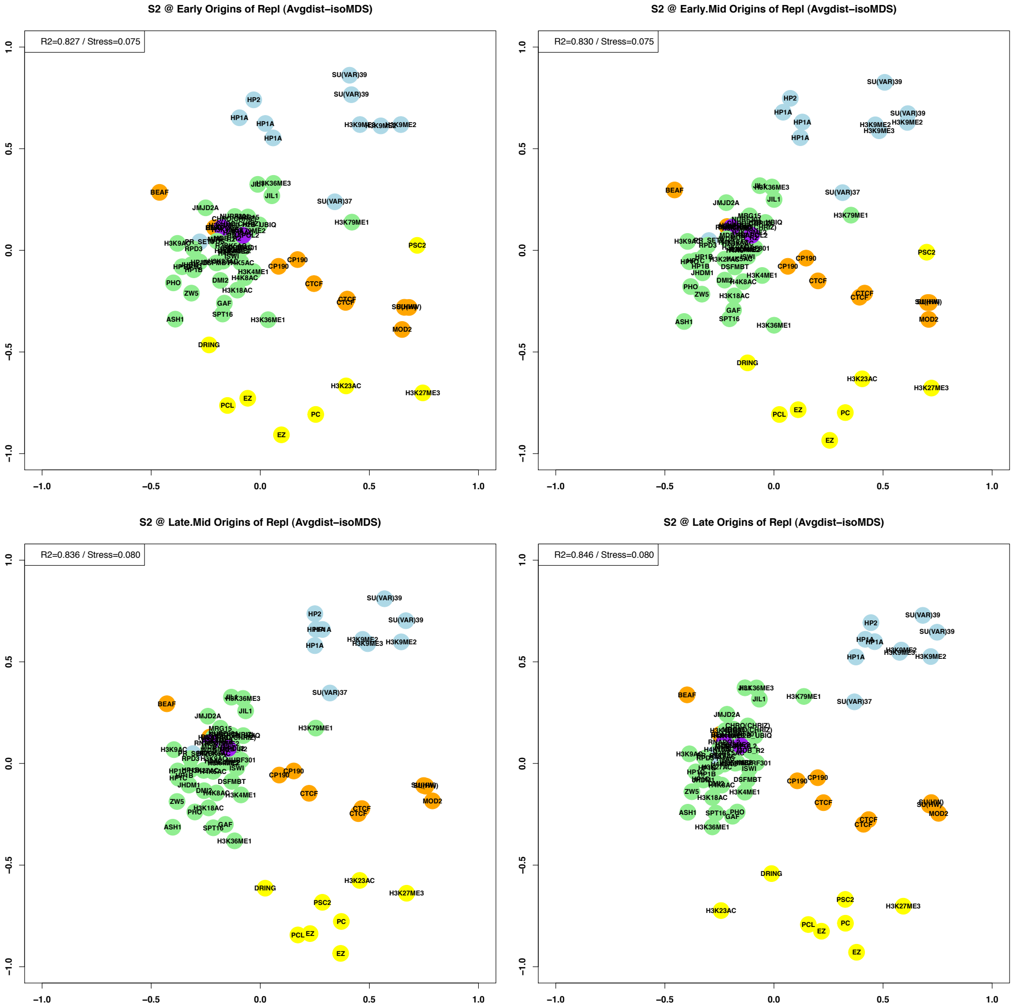

Supplementary Figure 5: chroGPS-factors map showing the epigenomic landscapes *Drosophila melanogaster* S2 cell lines at Early (top left), Early-Mid (top right), Mid-Late (bottom left) and Late (bottom right). Upon visual inspection we can already observe changes in location and distribution of some epigenetic factors and domains.

list of Procrustes errors for all common factors involved in the map. Statistical significance of the observed changes can be assessed via overlap permutation tests. Supplementary Figure 6 illustrates results for Procrustes differential analysis of the transition between Early-Mid and Late time points.

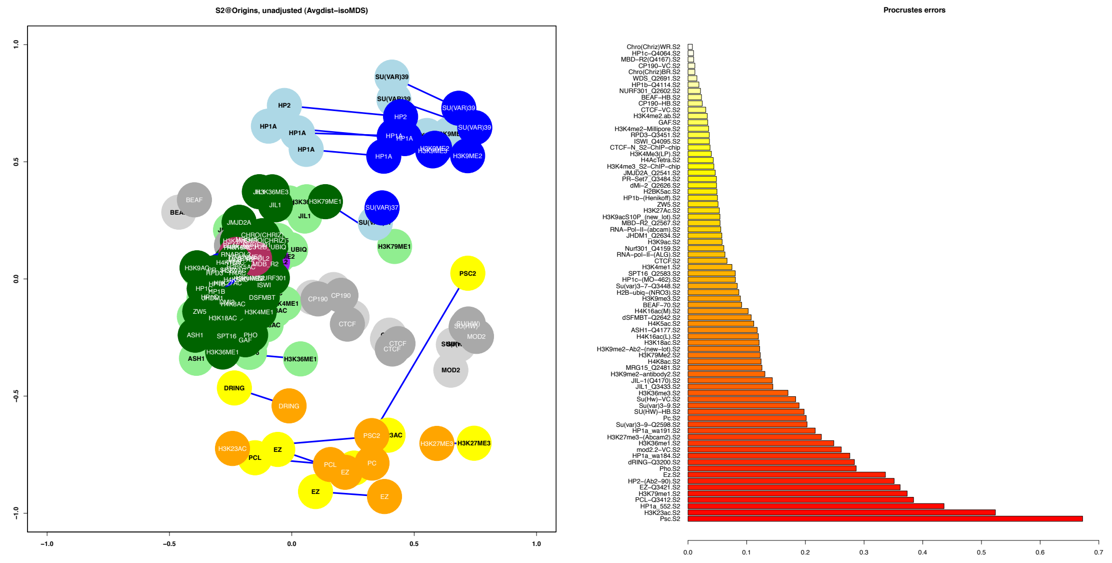

Supplementary Figure 6: Differential chroGPS-factors map of *Drosophila melanogaster* S2 cells at Early vs Late Origins of Replication (left), and Procrustes normalized errors (right). In this differential map, PSC2, a Polycomb related protein involved in non-heterochromatic gene silencing, is strongly shifted from the boundary-insulator region (Early) towards the Polycomb repression one (Late).

### 4 COMPARING CHROGPS-GENES MAPS

In chroGPS-genes maps, all genes in the genome are assigned a unique epigenetic 'profile', based on the epigenetic marks present at gene level in a certain biological scenario. Gene maps can also be used to highlight differences happening between different biological backgrounds such as different cells or tissues, diseases, and developmental or differentiation stages. In the following example we will explore several of these scenarios, using chroGPS to study differences between different epigenomic landscapes, as well as introduce the necessary methodology for performing this task.

#### 4.1 Generation and functional analysis of chroGPS-genes maps.

The procedure to generate and analyze chroGPS-genes maps is detailed in our previous publication [Font-Burgada et al., 2014] as well as in the chroGPS package vignette, however here we offer a summarized how-to highlighting the general steps.

The initial set-up for generation of chroGPS-genes maps is a collection of epigenetic factors in the shape of a *GenomicRanges* list, and a set of genetic elements (i.e. genes). chroGPS-genes are aimed at visualization and analysis of genetic elements based on their epigenetic state, that is, the epigenetic marks they present. In this example we will focus on genes (as defined by the longest possible transcript for each gene in the UCSC dm3 genome. The base object for generating chroGPS-genes maps is a matrix of $G$ rows (usually genes) and $F$ columns (epigenetic factors), and where each cell $Gi,Fj$ of the matrix will take a value of 1 if a certain epigenomic mark j is reported for gene i, and 0 otherwise.

The resultant binary matrix is given as the main argument to the *distGPS* function, that will compute pairwise similarities/distances between rows of the matrix (in our case, genes), based on their epigenetic profiles. Thus, genes sharing a high number of common factors will present a high similarity value (or small distance), whereas ones sharing very few ones will be the opposite. As with chroGPS-factors, we offer different similarity metrics and ways to weight and penalize presence and absences of epigenetic factors when computing distances, as well as different ways to deal with technical replicates of the same epigenetic factor.

The following step is to use the resulting distGPS object to generate a low-dimensional representation using MDS, as we did in chroGPS-factors. In order to speed up computation of the MDS solution with gene maps that comprehend thousands of genes, we offer parallel computation using a random split-and-combine approach that also offers optimization of the resulting map in order to maximize goodness-of-fit and removes potential undesired effects from the randomization applied in the parallel computation.

After generation of the map, the next natural step in our approach is to perform clustering analysis using the clusGPS function, which by default performs a hierarchical clustering of the original distance matrix using average linkage. Methods for unsupervised determination of underlying number of clusters in our scenario and Bayesian non-parametric density estimation for assigning posterior probabilities of cluster identity for each element in the map are provided [Jara et al., 2011]. Final map visualization can also incorporate additional useful information such as expression values by means of palette color scales in the provided plot function, and point size to reflect cluster classification uncertainty. Functional analysis of the identified clusters can also be performed by means of assessing their enrichment / depletion on epigenetic marks when compared to the whole map using the *profileClusters* function. Supplementary Figure 7 shows final result of this approach as analyzed in [Font-Burgada et al., 2014].

#### 4.2 Introduction to chroGPS-genes differential maps

Generation of differential chroGPS-genes maps to compare two epigenomic data sets is performed in a very similar way to the ones presented above. First, common factors between both backgrounds to compare are selected, usually after performing some operation to unify factor replicates when present, using the *mergeReplicates* and *combineGenesMatrix* functions (See Supplementary Figure 8). Then, for genes with at least one epigenetic mark in each background we compute pairwise distances between their epigenetic profiles using our metric of choice and a 2 dimensional map is generated via MDS. This allows to keep trace both of the epigenetic and background identity for each gene in the analysis. Since genes can be identified easily on the map by means of its epigenetic profile, it is straightforward to know which genes from different conditions present the same or very similar profiles (thus they are located in the same exact point or very close in the map), and which ones present significant differences (and therefore differ strongly in their location).

Downstream functional analysis of this differential map is essentially done in the same way as in a regular chroGPS-genes one. Briefly, hierarchical clustering is performed over the mathematical distances computed between each pair of unique epigenetic profiles. The identified clusters in this first step are then subject to an unsupervised merging process in order to further refine cluster definition and reduce granularity inherent to this method.

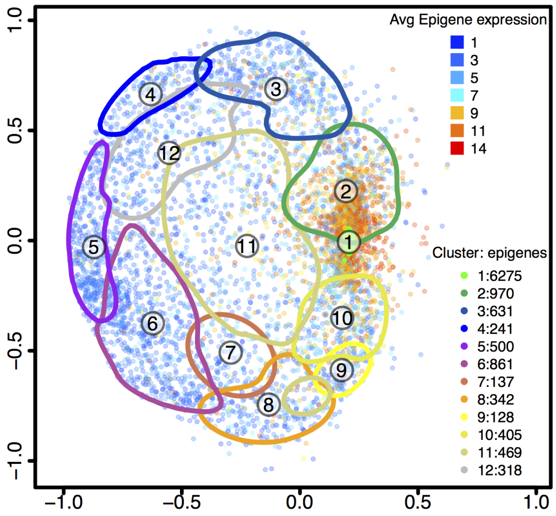

Supplementary Figure 7: chroGPS-genes map of S2 cells. The map is analyzed using hierarchical clustering with average linkage. Clusters corresponding to 50 percent between-cluster distance are shown after unsupervised merging. The epigenetic state of each cluster is determined based on the log2 enrichment/depletion ratio of each factor. 'Epigenes' are colored according to their average log2 RMA expression levels in S2 cells [Kessler et al., 2015], and they distribute clearly according to Active and Repression domains. Legends on the right side indicate number of genes in each cluster and average gene expression level of each gene point.

Finally, after a final clustering configuration is obtained, each gene is assigned a posterior probability of correct classification by means of Bayesian density estimation procedures, and the *diffGenes* function can be used to identify statistically significant changes epigenetic profiles at gene level.

#### 4.3 Comparing *Drosophila melanogaster* S2 and BG3 cell lines

We illustrate this with a case study comparing two well characterized *Drosophila melanogaster* cell lines, S2 and BG3. Schneider 2 cells, usually abbreviated asS2 cells, are one of the most commonly used Drosophila melanogaster cell lines. S2 cells were derived from a primary culture of late stage (20–24 hours old)*Drosophila melanogaster* embryos, likely from a macrophage-like lineage. BG3, however, is a cell line derived from central nervous system of third instar larvae.

To demonstrate how this strategy allows to identify changes related to biological differences between these two cell lines, following generation and functional analysis of the differential map involving common epigenetic factors present in S2 and BG3 we iterate over gene groups involved in cluster transitions between both backgrounds, and proceed to perform a Gene Ontology enrichment analysis for all genes involved in the observed cluster transitions.

The usual scenario in which to perform a differential chroGPS-genes map is that one involving studying epigenetic profiles over genes in two different conditions, biological or technical backgrounds from the same species, even though one could use the same methodology to address for instance changes in regulatory mechanisms in promoter vs coding regions or origins of replication, by simply recomputing binding site assignments under different genomic locations, etc. The basic elements to generate this map are the two collection of epigenetic elements which we want to compare at our desired level to use for generation of the necessary binary matrices described previously, or in its defect, two already computed matrices of 0s and 1s representing epigenetic profiles for each genetic element in each condition. Only common epigenetic factors mapped in both conditions will be taken into account (and therefore, only common genetic elements with at least 1 mapped factor in each condition are used). The *combineGenesMatrix* function takes raw *genes* x *factors* matrices for both backgrounds and returns a unique matrix containing unique epigenetic profiles for them (see Supplementary Figure 8).

| | | F1 | F2 | ... | Ff-1 | Ff |
|---|---|---|---|---|---|---|
| S1 | G1 | 1 | 0 | | 0 | 1 |
| | G2 | 0 | 0 | | 1 | 0 |
| | ... | ... | ... | ... | ... | ... |
| | Gn-1 | 0 | 1 | ... | 0 | 0 |
| | Gn | 0 | 0 | | 0 | 1 |
| S2 | G1 | 1 | 0 | | 0 | 1 |
| | G2 | 0 | 0 | | 1 | 1 |
| | ... | ... | ... | ... | ... | ... |
| | Gn-1 | 0 | 1 | ... | 0 | 0 |
| | Gn | 0 | 1 | | 0 | 1 |

| | F1 | F2 | ... | Ff-1 | Ff |
|---|---|---|---|---|---|
| EP1 | 1 | 0 | | 0 | 1 |
| EP2 | 0 | 0 | | 1 | 0 |
| EP3 | 0 | 0 | | 1 | 1 |
| ... | ... | ... | ... | ... | ... |
| EPm-2 | 0 | 1 | ... | 0 | 0 |
| EPm-1 | 0 | 0 | | 0 | 1 |
| EPm | 0 | 1 | | 0 | 1 |

| EP1 | EP2 | EP3 | ... | EPm-2 | EPm-1 | EPm |
|---|---|---|---|---|---|---|
| D 1,1 | D 1,2 | D 1,3 | ... | D 1,m-2 | D 1,m-1 | D 1,m |
| - | D 2,2 | D 2,3 | ... | D 2,m-2 | D 2,m-1 | D 2,m |
| - | - | D 1,3 | ... | D 3,m-2 | D 3,m-1 | D 3,m |
| - | - | - | ... | ... | ... | ... |
| - | - | - | - | D m-2,m-2 | D m-2,m-1 | D m-2,m |
| - | - | - | - | - | D m-1,m-1 | D m-1,m |
| - | - | - | - | - | - | D m,m |

Supplementary Figure 8: Left: stacked binary matrix with presence/absence information for common genes and epigenetic factors from backgrounds S1 and S2. Center: The unique epigenetic profiles for both backgrounds are used to compute a unique similarity/distance matrix based on pairwise similarity between the vectors for all epigenetic profiles (right).

This combined matrix is given as input to the *distGPS* function, together with the respective labels identifying each background. The resulting object is used to generate a low-dimensional map of all genetic elements based on the similarity of their epigenetic profiles using MDS. All other methodologies used with regular chroGPS-genes maps (clustering, etc) are available for differential maps as well. To sum up, the map just represents a certain number of genetic elements on space based on similarity of their epigenetic profiles. Thus, two genes sharing exactly the same epigenetic profile will be located at exactly the same point, whereas changes in their profiles will translate into shifted positions. These changes and their potential functional effects are indentified and analyzed with the *diffGenes* function.

This function takes as input the binary matrix with combined epigenetic profiles for both backgrounds and the clustering object produced by *clusGPS*, and retrieves cluster identities and posterior probabilities of classification. This information is provided for each genetic element analyzed and for both conditions of interest, and therefore it is straightforward not only to have a trace of all epigenetic cluster changes taking place for all analyzed genetic elements, but also to rank those changes based on confidence estimates as provided by the posterior probabilities of classification.

Once cluster transitions are obtained, a very straightforward approach for downstream analysis and is to perform Gene Ontology enrichment analysis via custom hypergeometric tests or using functions such as *getEnrichedGO* from the *ChIPpeakAnno* package, or using, DAVID, Gene Set Enrichment Analysis or others. In order to illustrate this, we show results of custom hypergeometric tests with Gene Ontology datasets to investigate potentially interesting Biological Process terms enriched among genes involved in cluster transitions.

Between all cluster transitions obtained (See Supplementary Table 1), in Supplementary Figure 9 we present the differential map highlighting results for genes involved in cluster transition 5 (S2) to 2 (BG3). Top Gene Ontology enriched results from the list of 70 genes involved in this cluster transition present a strong and statistically significant enrichment in processes such as axon guidance, neuron growth and other central nervous system ones.

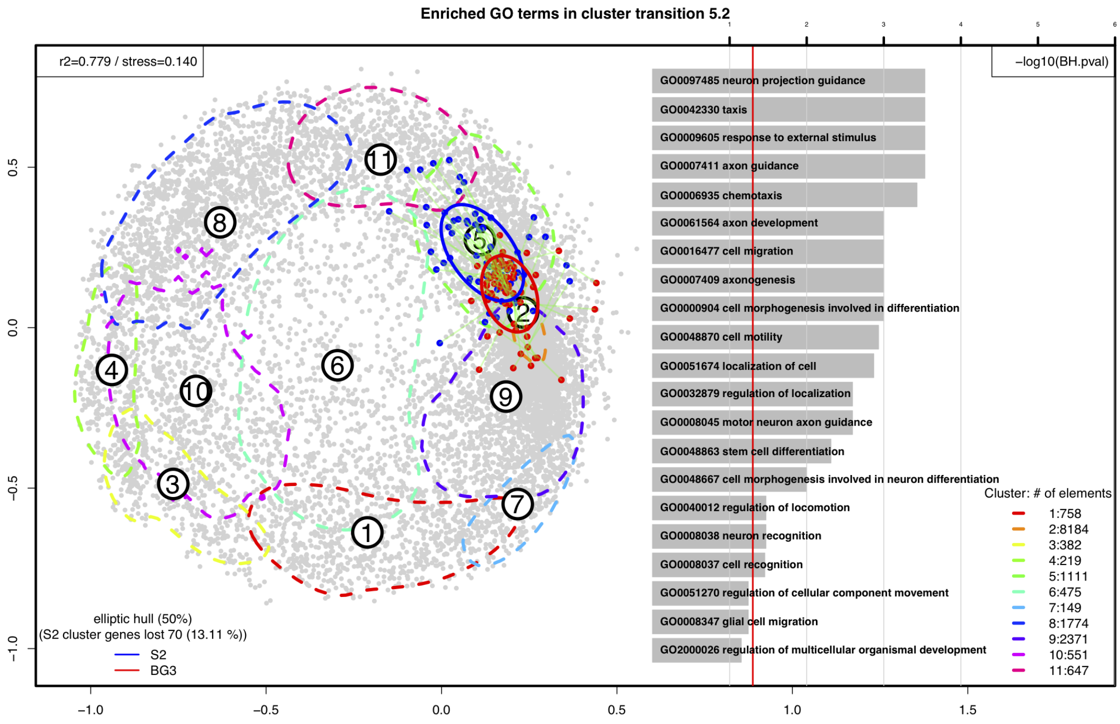

Supplementary Figure 9: chroGPS-genes differential map of *Drosophila melanogaster* S2 and BG3 cells showing top 20 Biological Process enriched terms for genes involved in cluster transition 2/5.

### 5 SOME NOTES ON VISUALIZATION

#### 5.1 Exporting chroGPS maps to XGMML (Cytoscape) and HTML5 (Plotly)

Maps generated with chroGPS can be exported to GraphML XGMML format using the provided funtion gps2xgmml. These maps can be imported with several R network analysis packages such as *igraph* and are also suitable to use with external network visualization and analysis solutions like Cytoscape. Additional information regarding distances between elements in the map and all other information can be easily exported into tabular format files and assigned to the imported network nodes using the standard procedures for each software.

Plotly, also known by its URL, Plot.ly, is an online data analytics and visualization tool that provides online graphing, analytics, and statistics tools as well as scientific graphing libraries for Python, R, MATLAB, and other languages. The function *getPoints* can be used to retrieve point coordinates defining our epigenomic factor and gene map objects, leading to straightforward visualization using 2 and 3-dimensional scatterplots, and all additional metadata and functional and differential analysis results can be easily exported in the form of plain tab-delimited text files to use in advanced and customized static or dynamic visualization solutions.

### 6 DATA AND CODE AVAILABILITY

The chroGPS2 package with routines for generation, visualization and funcional analysis of epigenome maps is available in Bioconductor. The package includes small toy example datasets, code snippets and a detailed user manual illustrating the main functionalities presented in this supplementary material. Complete datasets presented in this work available upon request.
